# Dynamin A as a one-component division machinery for synthetic cells

**DOI:** 10.1101/2022.12.05.519112

**Authors:** Nicola De Franceschi, Roman Barth, Sabrina Meindlhumer, Alessio Fragasso, Cees Dekker

## Abstract

Membrane abscission, the final cut of the last connection between emerging daughter cells, is an indispensable event in the last stage of cell division, as well as in other cellular processes such as endocytosis, virus release, or bacterial sporulation. However, its mechanism remains poorly understood, which also impedes its application as a cell-division machinery for synthetic cells. Here, we use fluorescence microscopy and Fluorescence Recovery After Photobleaching (FRAP) to study the *in vitro* reconstitution of the bacterial protein Dynamin A (DynA) inside liposomes. Upon external reshaping of the liposomes into dumbbells, DynA self-assembles at the membrane neck, resulting in membrane hemi-scission and even full scission. DynA proteins constitute a simple one-component division machinery that is capable of splitting dumbbell-shaped liposomes, marking an important step towards building a synthetic cell.

## Introduction

The ability to divide is one of the most fundamental features of cellular life. Since all cells are surrounded by a lipid membrane, the division process in all organisms necessarily entails a significant membrane deformation. In a first step, the mid-cell region of the cell is constricted, resulting in the formation of a so-called dumbbell-like shape, where the two future daughter cells are still connected by a narrow neck. In eukaryotes, this process is driven by an actomyosin ring underneath the membrane^1^, whereas constriction in bacteria is coordinated intracellularly by the Z-ring^2^ with peptidoglycan synthesis occurring at the outside of the membrane^3^. Following constriction into the dumbbell geometry, the final step of cell division consists of the membrane scission of the neck (also termed abscission). This is believed to be performed by yet other protein machineries such as the ESCRT-III complex in eukaryotes^4^ and the evolutionary related Cdv complex in Archaea^5^, while it is not yet fully clear how abscission is accomplished in bacteria.

Membrane remodelling has been extensively studied in reconstituted systems ^6^. Membrane abscission can be studied in various membrane geometries, i.e., where proteins act from the *outside* on a membrane neck, or in a geometry where a membrane-scission machinery acts from the *inside* of a membrane neck, the so-called “reverse topology” (Figure S1) ^7^. A classic example of the former is endocytosis, where eukaryotic dynamin binds to the outside of a membrane neck and induces scission of endocytic pits^8^. Reverse-topology processes include scission events mediated by the ESCRT-III complex in eukaryotes^9^, and cell division and sporulation in bacteria^10^. Recent years witnessed a growing interest in building a self-sustaining and self-reproducing synthetic cell^9^ – a liposome filled with proteins that exhibits features of natural cells. Cell division of such synthetic cells should occur in reverse topology as components of the division machinery need to be synthesized within the synthetic cell. However, such a division system has so far been missing ^11^.

In the filamentous bacteria *Streptomyces*, two dynamin-like GTPases, DynA and DynB, localize near a septum where they induce the final stage of sporulation^12^. While the exact function of DynA/B in *Streptomyces* remains incompletely understood, bacterial dynamin-like proteins have been widely associated with membrane remodelling events^13^. For example, *B. subtilis* DynA, which is a fusion protein that combines the functionalities of both DynA and DynB within one protein^14^, has been shown to mediate Mg^++^-dependent membrane fusion *in vitro* by binding to the headgroups of negatively charged lipids, while GTP hydrolysis was found to be dispensable for this process^15^. Moreover, DynA was shown to counteract stress-induced pores ^16^ during phage infections ^17^. Notably, membrane scission and fusion involve the same “topological reaction”, but run in opposite directions – from one to two vesicles, or vice versa (Figure 1a).

**Figure 1:**
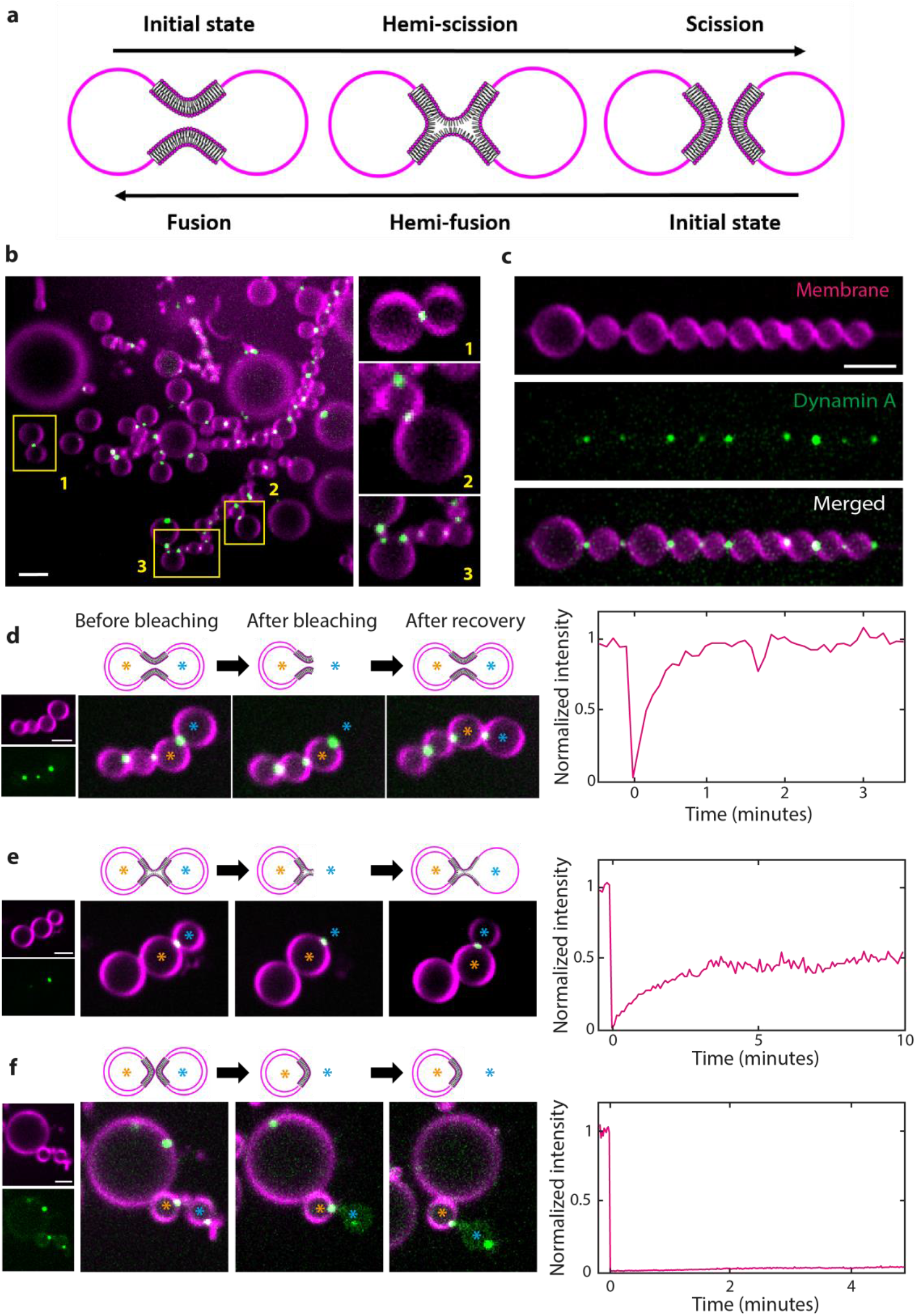
Dynamin A localizes at necks of dumbbells and affects membrane connectivity. **(a)** Schematics depicting the fusion/scission “topological reaction”. **(b)** Large field of view of a preparation of dumbbells with encapsulated DynA generated with the SMS. Insets show detailed views of DynA clusters localized at membrane necks. **(c)** Example of a chain of dumbbells with DynA clusters at multiple necks. **(d)** Full recovery of fluorescent lipids to a normalized intensity N_I_≈1 upon photo-bleaching of one lobe of a chain of dumbbells. The bleached and the control lobes are indicated by a blue and an orange asterisks, respectively. The right panel shows the Normalized Intensity (N_I_) versus time. Schematics on the top illustrate the membrane connectivity at the neck that is compatible with the recovery profile. **(e)** Partial recovery of fluorescent lipids to a normalized intensity N_I_≈0.5 upon photo-bleaching of a lobe of a chain of dumbbells. Colours, time trace, and schematics are as in panel a. **(f)** Absence of recovery of fluorescent lipids upon photo-bleaching of a lobe of a chain of dumbbells. Colours, time trace, and schematics are as in panel a. The gain in the DynA fluorescence channel was increased to show the continuous presence of the bleached lobe. All scale bars: 5μm.

Inevitably, they both also entail the same intermediate: a hemi-scission/hemi-fusion state, where the split inner leaflet defines two distinct systems, while the outer leaflet still joins both vesicles (Figure 1a).

Here, we address the question whether bacterial *B. subtilis* DynA protein is able to mediate membrane scission when reconstituted *inside* membrane necks. We reconstituted recombinant DynA inside liposomes that were deformed into a dumbbell shape using the recently developed Synthetic Membrane Shaper (SMS) approach^18^. Using fluorescence microscopy and Fluorescence Recovery After Photobleaching (FRAP), we monitored whether soluble dyes within the liposomes and fluorescently labelled lipids in the bilayer leaflets could diffuse across the membrane neck. The experiments revealed that DynA localizes at the dumbbell neck where it catalyses membrane hemi-scission, which subsequently progresses to full scission. Our data demonstrate a novel function for DynA and provide insights into the process of membrane scission in reverse topology, which is the biologically relevant geometry for cell division. Moreover, this dynamin-based single-protein system presents an elegant novel synthetic divisome that is of interest for establishing division in synthetic cells^11^.

## Results

A high yield of dumbbell-shaped liposomes was obtained using the SMS technology that we recently introduced^18^. In this approach, droplets were generated by pipetting an “inner” aqueous solution into a lipid-in-oil suspension that was subsequently placed on top of a water reservoir, referred to as the “outer” buffer. Liposomes were then formed when the droplets crossed the oil-to-water interface (between oil and outer buffer) as they slowly sank due to gravity. The presence of small (96.5kDa) DNA structures called nanostars (which basically are Holliday Junctions armed with cholesterol moieties; hereafter collectively referred to as “CN”), in the outer solution caused their binding to the outer membrane during liposome formation, inducing membrane curvature, which led to a high yield of dumbbell-shaped liposomes.

When DynA was included in the inner solution, a strong preferential localization of DynA was observed at the membrane necks in the resulting dumbbell structures (Figure 1b), as evidenced by the presence of a bright fluorescent spot at the neck (Movie 1), whereas binding of DynA to the liposome membrane outside the neck region was minimal. The fact that DynA strongly localizes at the necks indicates that it prefers to bind to highly curved membranes. For the present work, we chose to focus on chains of dumbbells (Figure 1c), which constituted a sizable fraction of the liposomal structures in our samples. Such chains of dumbbells, which consist of a linear array of many liposomes that are mutually connected by a neck, result from a shape transformation of a continuous membrane system during the process of liposome formation (Movie 2). The chain topology, where both leaflets of the membrane are continuously connected, presents a useful framework to study the function of DynA at dumbbell necks in reverse topology.

In order to probe the nature of the membrane connectivity between lobes, we performed FRAP experiments, where the fluorescent lipids of one liposome within the chain were photobleached. The fluorescence intensity subsequently recovered over time as lipids flowed across the neck region that connected the adjacent lobes^18^, see Figure 1d-f. We quantified the degree of fluorescence recovery by the normalized intensity (N_I_), which measures the ratio between the final fluorescence intensity of the bleached lobe after recovery and that of a neighbouring control lobe that was not photobleached (for details, see Supplementary Note 1, Figure S2, S3). This intensity N_I_ is normalized such that a final value of N_I_=1 indicates full recovery, while N_I_=0 signifies the absence of any recovery.

We observed three distinct outcomes of these FRAP experiments on liposome chains that were otherwise indistinguishable from each other as they all were connected by necks harbouring a DynA cluster. In some chains, full recovery occurred to N_I_=1 (Fig.1d; Movie 3), indicating that both leaflets of the membranes of the two adjacent lobes were fully connected and supported lipid diffusion. This indicates the presence of an open neck, as also confirmed from control FRAP experiments on dumbbell-shaped liposomes without DynA, which yielded the same result (Figure S4, S6b). A second set of data showed, even after prolonged time, only a partial recovery that plateaued at a N_I_ value somewhere between 0 and 1 (Fig.1e; Movie 4). This is consistent with a scenario where hemi-scission had occurred, meaning that the inner leaflet of the membrane had undergone fusion (thus preventing lipid diffusion across the neck), while lipids in the outer leaflet could still freely flow between the lobes, yielding a partial recovery of the bilayer fluorescence signal. Finally, some dumbbells displayed no recovery at all in FRAP experiments (Figure 1f, Movie 5), consistent with a scenario where both leaflets had undergone scission. Even upon full scission, we observed that the lobes did not detach from each other, most likely due to the presence of Mg^++^ ions in the outer buffer which bridges negatively charged lipids in the membrane. However, at least in one case, we observed lobe detachment upon scission (Figure S5).

To quantify these data and to discern to what extent these membrane (hemi-)scission events can be attributed to the action of DynA, we performed lipid FRAP experiments on a large dataset of chains of dumbbells (n = 207) that were generated by having either (1) only bare membrane (BM), or (2) chol-oligo + nanostars on the outside (BM+CN), or (3) CN on the outside as well as DynA on the inside (BM+CN+DynA). Pooling all data together yielded a plot for the normalized intensity N_I_ with three distinguishable populations that we fitted with Gaussians (Figure S6a). From fitting, we extracted mean values for the three populations that correspond to no lipid recovery (average N_I,1=0.04_, close to 0), partial recovery (average N_I,2_=0.31), and full recovery (average N_I,3_=0.92, close to 1). Note that the N_I,2_=0.31 peak is very close to the expected value of 0.33 for an ideal case of membrane hemi-scission (Supplementary Note 1, Figure S2). Identification of these peaks with events of, respectively, no scission (open neck), hemi-scission, and full-scission events, allowed us to quantify the number of these events in the different profiles for N_I_ for each experimental condition.

As Figure 2a shows, the vast majority (85%) of dumbbell liposomes formed by the nanostars featured an open neck, as only a marginal increase of membrane hemi-scission (8%) and full scission (7%) events was observed, compared to dumbbells having only bare membranes which showed virtually all open necks (Figure S6b). This, however, strikingly changed upon the addition of DynA. Co-reconstitution of both DynA on the inside and CN on the outside of chains of dumbbells resulted in 36% of necks in a hemi-scission state, and 26% in a full scission state (Figure 2b). This shows that DynA is able to induce both membrane hemi-scission and full scission in these conditions. DynA was furthermore found to narrow down the neck diameter, as measured for the subset of dumbbells where neither hemi-scission nor full scission occurred (Figure S7). We estimated the degree of DynA enrichment at necks by calculating the Recruitment Ratio (RR), which compares the DynA fluorescence at necks with the fluorescence of residual protein present in the lumen of the liposome (see methods). Interestingly, we observed a correlation between the degree of DynA enrichment and the propensity of the necks to undergo membrane rearrangement (Figure 2c). Full scission occurred more frequently when the degree of enrichment was relatively low (RR = 8 ± 3, N=14, mean ± SD), while the neck was more likely to be trapped in a hemi-scission intermediate when a higher amount of DynA was assembled to the neck (RR = 28 ± 20, N=12, mean ± SD). This may indicate that excessive recruitment of DynA can impair the transition from hemi-scission to full scission.

**Figure 2:**
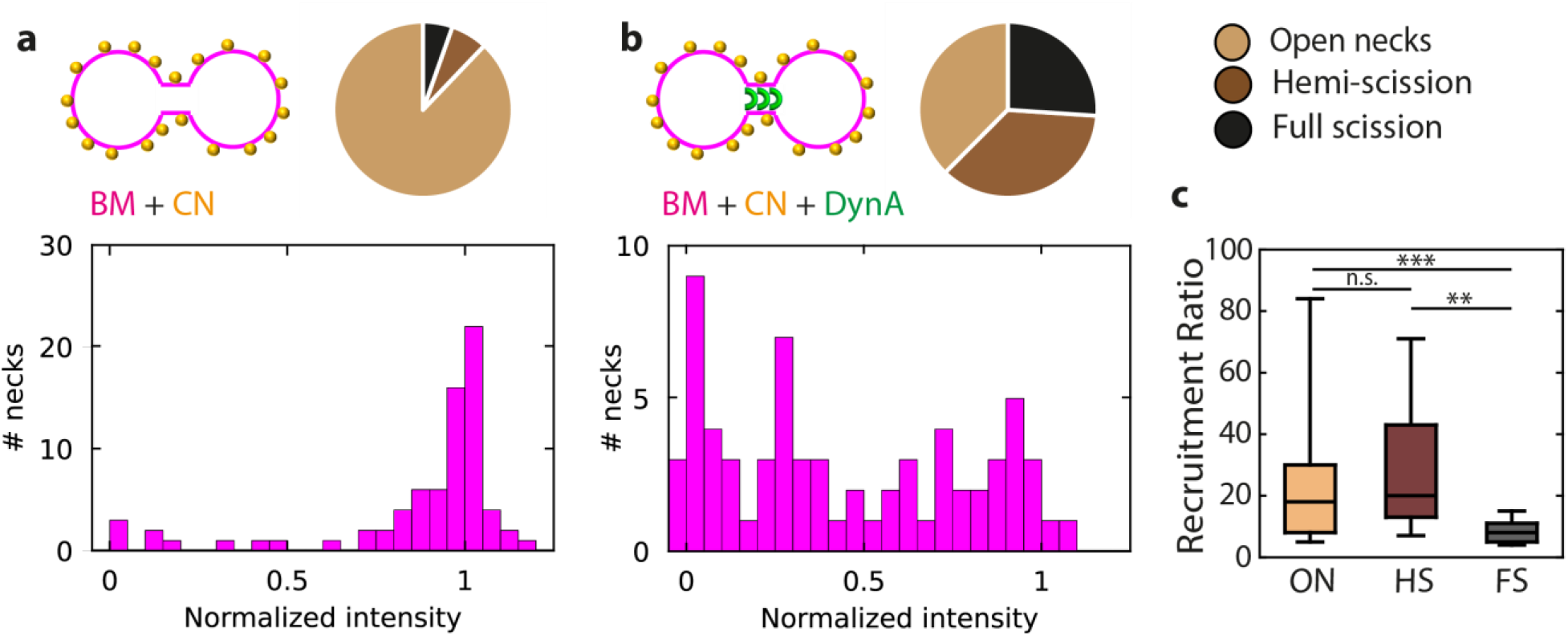
Quantification of membrane remodelling events at necks in the presence or absence of DynA. **(a)** Histogram of normalized intensities N_I_ for dumbbells with cholesterol-oligo and nanostars but without DynA (BM + CN; 75 necks from 7 independent preparations). Top-right: pie-chart indicating the fraction of open necks, hemi-scission, and full scission events. BM = bare membrane; CN = cholesterol-oligos + Nanostars. **(b)** Histogram of normalized intensities N_I_ for dumbbells with cholesterol-oligo, nanostars and DynA (BM + CN+DynA; 69 necks from 12 independent preparations). Top-right: pie-chart indicating the fraction of open necks, hemi-scission, and full scission events. **(c)** Recruitment ratio RR of DynA. ON = open necks; HS = hemi-scission; FS = full scission. n=18 for open necks; n=12 for hemi-scission; n=14 for full scission. p-value=0.35 for open necks versus hemi-scission; p-value=0.006 for open necks versus full scission; p-value=0.0004 for hemi-scission versus full scission using the Mann-Whitney U Test.

In order to verify that the partial lipid recovery in FRAP experiments indeed corresponds to membrane hemi-scission, we encapsulated a soluble dye and performed simultaneous photobleaching of the lipids in the membrane and of the dye that was encapsulated in the lumen of the liposome. In the absence of DynA localization at the neck, dumbbells always exhibited full recovery of both the lipids and of the soluble dye (Figure 3a). This indicates that lipids and the soluble dye could freely diffuse across the necks, as expected. In the presence of DynA, however, we often observed that upon bleaching of both dye and lipids, the lipids recovered only partially (as described above, cf. Figure 1d), while the soluble dye in the inner volume of the liposome did not recover at all (Figure 3b). This corroborates the presence of a hemi-scission state, which allows lipids from the outer leaflet to recover, while fully preventing both the lipids of the inner leaflet and the soluble dye from flowing across the neck.

**Figure 3:**
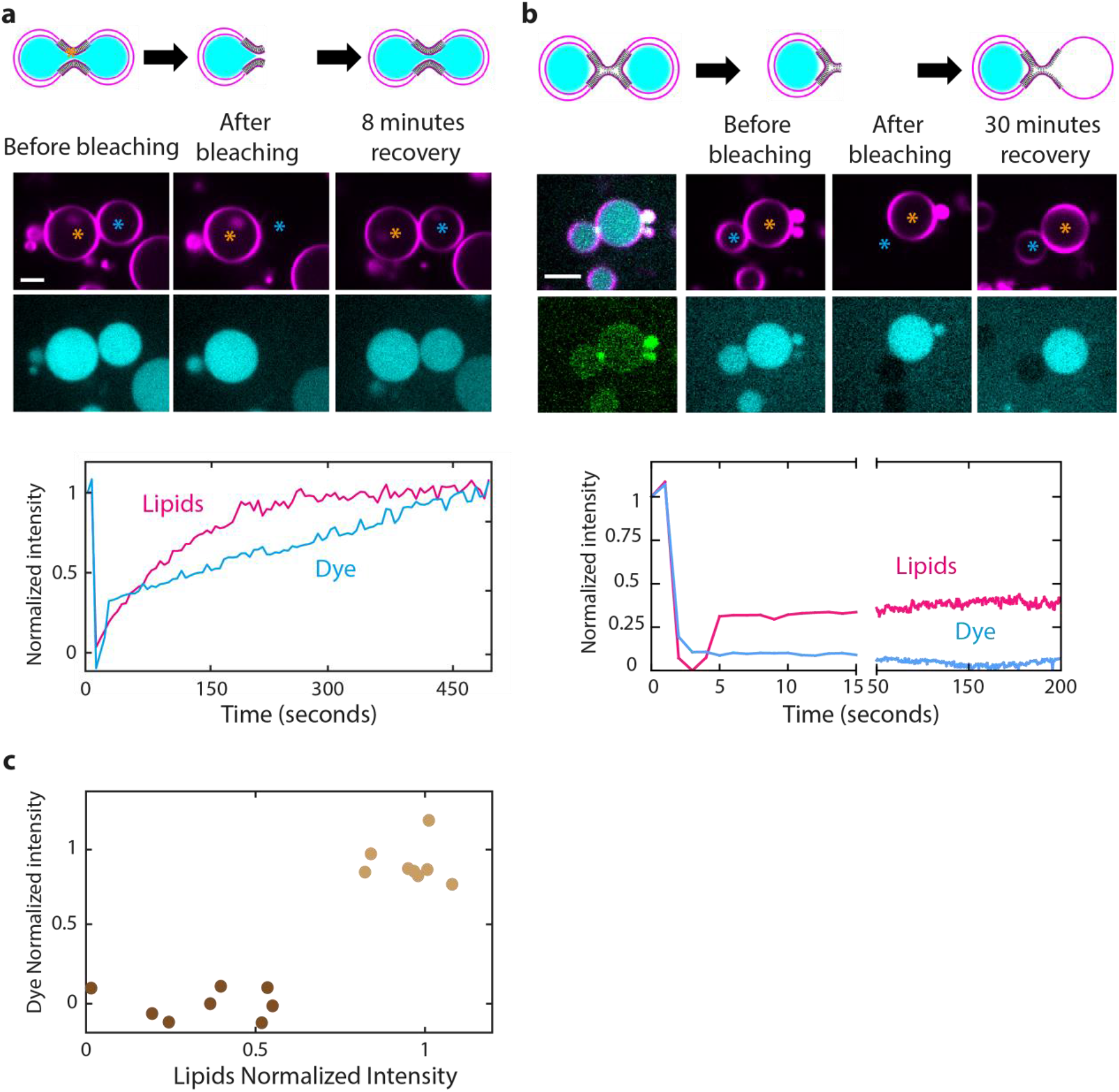
Characterization of DynA-induced membrane hemi-scission events. **(a)** Concomitant FRAP of lipids and soluble dye in dumbbells in the absence of DynA. Bottom: Normalized Intensity versus time of both lipids and dye. **(b)** Same as panel a but in the presence of DynA **(c)** Scatter plot showing the fraction of open necks, hemi-scission, and full scission events based on lipid recovery and the correspondent recovery or absence of recovery of the soluble dye (n=16). Open necks are indicated by in light brown; hemi-scission and scission events in dark brown.

We quantified the degree of recovery of both dye and lipids for each dumbbell, yielding the scatter plot shown in Figure 3c. In about half of the dumbbells, we observed a full recovery of both dye and lipids, indicating the presence of an open neck. In the remaining 50%, however, we observed either a partial lipid recovery or a total absence of lipid recovery, consistent with the hemi-scission and full scission data shown in Figure 2. Importantly, in 100% of cases in which the soluble dye did not recover, we observed either partial or a complete absence of lipid recovery. This correlation is expected for a closed neck, which prevents the dye from recovering. This can happen either by full scission, in which case no lipid recovery is observed, or by hemi-scission, where scission of the inner leaflet would allow partial lipid recovery. The data thus confirm that partial lipid recovery corresponds to a hemi-scission state.

## Discussion

In this paper, we showed that bacterial Dynamin A is able to trigger membrane scission when assembled inside membrane necks in dumbbell-shaped liposomes. With this demonstrated functionality, Dynamin A adds to the family of membrane-remodelling proteins involved in fusion and scission (Figure 4a). In earlier experiments, eukaryotic dynamin was shown to be able to induce membrane scission when assembled on the outside of membrane nanotubes^19^ and a hemi-scission intermediate was visualized^20^. The present work is the first example of visualization of membrane hemi-scission triggered by a protein assembled inside membrane necks.

**Figure 4:**
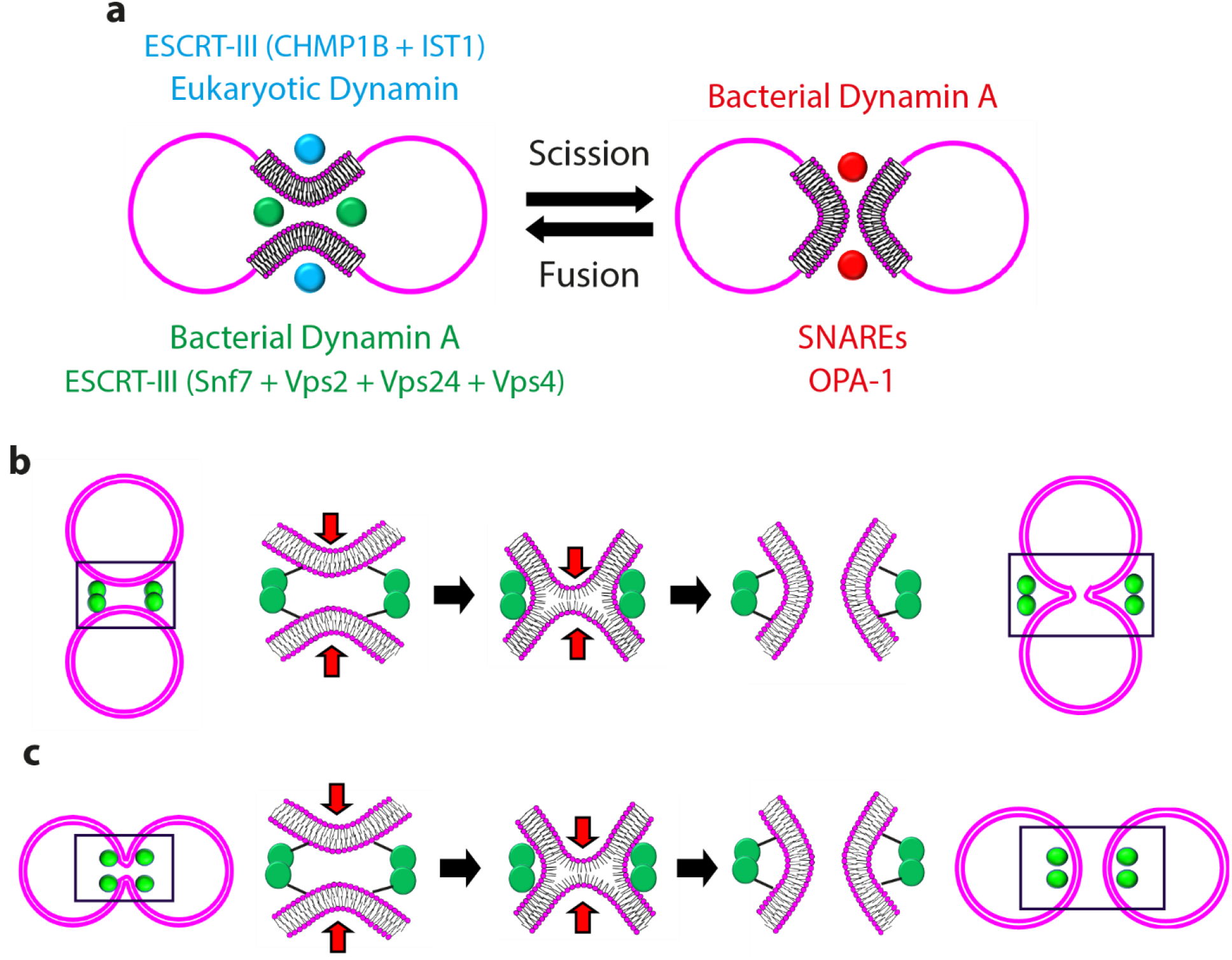
DynA in membrane scission and fusion. **(a)** Schematics summarizing the proteins involved in membrane scission and fusion. The dots indicate the position of the proteins with respect to the membrane and colours correspond to the protein names listed. The membrane remodelling activity of all these proteins has been confirmed by previous *in vitro* reconstitutions^15,29,34–37^, while the scission activity of bacterial DynA is established here. **(b)** Schematics representing membrane fusion via a hemi-fusion intermediate with DynA (in green) reconstituted on the *outer* leaflet. Red arrows indicate the direction of the membrane deformation induced by DynA. **(c)** Schematics representing membrane scission via a hemi-scission intermediate with DynA (in green) reconstituted on the *inner* leaflet.

Bacterial DynA was previously shown to promote membrane fusion when assembled on the outer leaflet of liposomes *in vitro*^14^. Our work is the first demonstration of membrane *scission* activity by a bacterial member of the Dynamin superfamily and, to the best of our knowledge, bacterial DynA is the only example of a protein able to trigger both membrane fusion and scission. DynA was previously modelled as a protein that can tether membranes in trans^13^, a configuration that – due to the symmetry of this topological reaction – can indeed explain liposome fusion as well as scission when DynA is present at the outside or inside, respectively (Figure 4b,c).

Previous attempts to induce membrane scission in dumbbell systems relied on laser irradiation^21^ or curvature induced by proteins that were bound to the outer leaflet^22^. In these reports, the occurrence of membrane scission was primarily established by observing the separation of daughter liposomes. It is, however, very common for liposomes to remain connected by lipid nanotubes, but such nanotubes are notoriously difficult to detect due to their intrinsic low fluorescence intensity (cf. Figure S8). Instead of apparent vesicle separation, we therefore settled to analyse our dumbbells using FRAP, which is the more rigorous way to establish membrane connectivity. Furthermore, we performed our experiments on linear chains of dumbbells, which originated from a single membrane system (Movie 2). Such long linear chains are unlikely to be the result of liposome adhesion, which would rather yield random aggregations.

Interestingly, we observed that about half of the membrane remodelling events triggered by DynA arrested at the hemi-scission stage (Figure 2b, 3b), yielding intermediates that are stable enough to be imaged for prolonged periods of time. While DynA has the capability to realise full scission, we often found it to do so only partially. This may be because DynA binds to the lipid headgroups on the inner leaflet of the membrane neck, while it does not directly interact with the outer leaflet (Figure 4c). Several factors may play a role in the progression from hemi-scission to full scission. This transition can occur spontaneously when the neck is thin enough, as it has been estimated *in silico* for necks with an inner radius of about 3 nm^23,24^, and in such a scenario, DynA’s role may merely be to reduce the neck width to such low diameters. Furthermore, the transition to full scission is favoured by membrane tension^25^, and our setup with the SMS system induces a dumbbell shape to the liposomes with a finite membrane tension^18^ that may act to assist the (hemi-)scission action of DynA. Finally, we observed that the transition to full scission is hindered by excessive accumulation of DynA at the neck (Figure 2c).

The only other protein machinery that so far was demonstrated to mediate membrane scission in an inverted topology is the eukaryotic ESCRT-III complex. This complex assembles inside membrane necks^26–28^ and triggers full membrane scission^29^ in *in vitro* tube-pulling assays. Notably, a hemi-scission intermediate was never visualized^4,30^. However, in these ESCRT experiments, the nanotube was kept under a pulling tension, which facilitated scission, while potential hemi-scission intermediates likely became short-lived and hard to detect. The alternative methodology we have introduced based on FRAP analysis of dumbbell liposomes allows to discriminate between different membrane connectivity states including hemi-scission. *In vivo*, it has been shown that the ESCRT-III complex completely disappears before full scission occurs^31^, suggesting that, similar to DynA, the action of the ESCRT-III complex may only induce hemi-scission.

Summing up, we showed that *B. subtilis* DynA promotes membrane scission in reverse topology, unveiling a new role for this protein. Due to its simplicity, this system presents an attractive candidate to build a divisome for synthetic cells^11^. A mainstream approach to build a synthetic cell relies on the use of reconstituted transcription/translation systems (RTTSs) to produce the proteins for growth and division^32^. The necessity of having multicomponent systems with defined stoichiometries complicates their implementation, however. Hence, a divisome having fewer components is obviously to be favoured. In this regard, a single-component system like DynA appears to be an excellent choice for building a synthetic divisome, as it presents advantages over more complex multi-component systems such as ESCRT-III.

## Methods

### Reagents

Glucose (G7021), MgCl_2_ (M8266), silicone oil (317667), and mineral oil (M3516-1L) and Optiprep (60% (w/v) iodixanol in water, D1556) were purchased from Sigma-Aldrich. Tris-HCl (10812846001) was purchased from Roche. DOPC (1,2-dioleoyl-*sn*-glycero-3-phosphocholine) (850375), DOPE-PEG(2000) Amine (1,2-dioleoyl-sn-glycero-3-phosphoethanolamine-N-[amino(polyethylene glycol)-2000] (ammonium salt)) (880234), 18:1 (Δ9-Cis) PG (1,2-dioleoyl-sn-glycero-3-phospho-(1’-rac-glycerol) (sodium salt)) (840475) and DOPE-Rhodamine (1,2-dioleoyl-*sn*-glycero-3-phosphoethanolamine-*N*-(lissamine rhodamine B sulfonyl) (ammonium salt) (810150C) were purchased from Avanti Lipids. Lipids were stored and resuspended in anhydrous chloroform (288306, Sigma Alrich). UltraPure bovine serum albumin used for passivation of the glass coverslips was purchased by ThermoFisher. For the FRAP experiments on soluble dye, Alexa Fluor™ 488 C5 Maleimide was used (A10254, ThermoFisher). Composition of solutions used in SMS preparations are shown in Supplementary Table 1 (inner solution) and Supplementary Table 2 (outer solution).

### DNA constructs

The 4 DNA oligos composing the cross-shaped nanostars were purchased from IDT. The sequence of the DNA oligos composing the nanostars and chol-oligo are reported in^18^.

### Protein purification and labelling

Bacillus subtilis DynA was overexpressed from pET16b (kindly provided by Dr. M. Bramkamp) and purified essentially as described^15^, but eluted with a linear imidazole gradient instead of a step elution. A solution of ~10 μM Dynamin was labelled with eightfold molar excess of Alexa Fluor 488 maleimide in the presence of 0.05 mM TCEP (45 minutes at room temperature), quenched with 10 mM β-mercaptoethanol and separated from free label on a Superdex S200 column equilibrated with T5 buffer (50 mM Tris/HCl pH8.0, 500 mM NaCl, 10 % glycerol).

### Lipid-in-oil suspension and droplets preparation

Lipid-in-oil suspensions were prepared according to^18^. Briefly, lipids solubilized in chloroform were mixed and blow-dried. They were re-solubilized with chloroform inside a glovebox, and a mixture of silicone and mineral oil^33^ was added. The resulting suspension was sonicated in ice for 15 min. The lipids mix used in this study was composed of DOPC (89.75% mol/mol), DSPE-PEG2000-biotin (2% mol/mol), DOPG (8% mol/mol) and DOPE-Rhodamine (0.25% mol/mol).

### Dumbbell preparation

The inner buffer was composed of 50mM Tris pH7.4 + 37% optiprep. The outer buffer was composed of 50mM Tris pH7.4 + 5mM MgCl2, to which glucose was added until reaching an osmolarity 40mOsm higher than the inner buffer. 10μM of each of the four oligomers composing the nanostars and 20μM of cholesterol-oligo^18^ were added to the outer buffer. DynA at 100nM was added to the inner buffer, droplets were generated by manual pipetting, and they were added to the outer buffer. For the experiment showed in Figure S5c, a DynA concentration of 20nM was used in order to minimize aggregation and to obtain dumbbells^18^.

### Data collection and analysis

Fluorescence images were acquired at the midplane of liposomes using spinning disk confocal laser microscopy (Olympus IX81 microscope, 60× objective, iXon camera) with Andor iQ3 software. To induce photobleaching, we employed raster scanning with a 491 nm laser (at 9.8 mW) over the region of interest. To measure the recovery signal, frames were collected every 1 s, starting right after the photobleaching event. Fluorescence images were analyzed and processed using ImageJ (v2.1.0). The extracted fluorescence data were plotted and fitted using Phyton 3. A detailed explanation of the image analysis pipeline is provided in Supplementary Note 1.

### Calculation of Recruitment Ratio

The Recruitment Ratio R_R_ at membrane necks was calculated with the following formula:

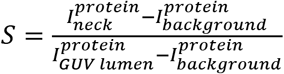

where 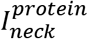 and 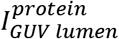 represent the fluorescence intensities of protein at the neck and of residual neck GUV lumen protein present in the lumen of the liposome, respectively.

## Acknowledgments

We thank S.J. Marrink and W. Pezeshkian for useful discussions, and M. Bramkamp for kindly providing the plasmid for the *B. subtilis* DynA. We acknowledge funding support from the BaSyC program of NWO-OCW and from ERC Advanced Grant 883684.

## References

1. Spira, F. et al. Cytokinesis in vertebrate cells initiates by contraction of an equatorial actomyosin network composed of randomly oriented filaments. Elife 6, (2017).

2. Allard, J. F. & Cytrynbaum, E. N. Force generation by a dynamic Z-ring in Escherichia coli cell division. Proc Natl Acad Sci U S A 106, 145–50 (2009).

3. Bisson-Filho, A. W. et al. Treadmilling by FtsZ filaments drives peptidoglycan synthesis and bacterial cell division. Science 355, 739–743 (2017).

4. Pfitzner, A.-K., Moser von Filseck, J. & Roux, A. Principles of membrane remodeling by dynamic ESCRT-III polymers. Trends Cell Biol 31, 856–868 (2021).

5. Caspi, Y. & Dekker, C. Dividing the Archaeal Way: The Ancient Cdv Cell-Division Machinery. Front Microbiol 9, 174 (2018).

6. Bassereau, P. et al. The 2018 biomembrane curvature and remodeling roadmap. J Phys D Appl Phys 51, (2018).

7. Hurley, J. H. ESCRTs are everywhere. EMBO J 34, 2398–407 (2015).

8. Sundborger, A. C. & Hinshaw, J. E. Regulating dynamin dynamics during endocytosis. F1000Prime Rep 6, 85 (2014).

9. Lemus, L. & Goder, V. Membrane trafficking: ESCRTs act here, there, and everywhere. Curr Biol 32, R292–R294 (2022).

10. Bohuszewicz, O., Liu, J. & Low, H. H. Membrane remodelling in bacteria. J Struct Biol 196, 3–14 (2016).

11. Olivi, L. et al. Towards a synthetic cell cycle. Nat Commun 12, 4531 (2021).

12. Schlimpert, S. et al. Two dynamin-like proteins stabilize FtsZ rings during *Streptomyces* sporulation. Proceedings of the National Academy of Sciences 114, (2017).

13. Bramkamp, M. Structure and function of bacterial dynamin-like proteins. Biol Chem 393, 1203–14 (2012).

14. Guo, L. & Bramkamp, M. Bacterial dynamin-like protein DynA mediates lipid and content mixing. FASEB J 33, 11746–11757 (2019).

15. Bürmann, F., Ebert, N., van Baarle, S. & Bramkamp, M. A bacterial dynamin-like protein mediating nucleotide-independent membrane fusion. Mol Microbiol 79, 1294–304 (2011).

16. Sawant, P., Eissenberger, K., Karier, L., Mascher, T. & Bramkamp, M. A dynamin-like protein involved in bacterial cell membrane surveillance under environmental stress. Environ Microbiol 18, 2705–20 (2016).

17. Guo, L., Sattler, L., Shafqat, S., Graumann, P. L. & Bramkamp, M. A Bacterial Dynamin-Like Protein Confers a Novel Phage Resistance Strategy on the Population Level in Bacillus subtilis. mBio 13, e0375321 (2022).

18. de Franceschi, N. et al. A synthetic membrane shaper for controlled liposome deformation. bioRxiv 2021.12.22.473854.

19. Antonny, B. et al. Membrane fission by dynamin: what we know and what we need to know. EMBO J 35, 2270–2284 (2016).

20. Mattila, J.-P. et al. A hemi-fission intermediate links two mechanistically distinct stages of membrane fission. Nature 524, 109–113 (2015).

21. Dreher, Y., Jahnke, K., Schröter, M. & Göpfrich, K. Light-Triggered Cargo Loading and Division of DNA-Containing Giant Unilamellar Lipid Vesicles. Nano Lett 21, 5952–5957 (2021).

22. Steinkühler, J. et al. Controlled division of cell-sized vesicles by low densities of membrane-bound proteins. Nat Commun 11, 905 (2020).

23. Kozlovsky, Y. & Kozlov, M. M. Membrane fission: model for intermediate structures. Biophys J 85, 85–96 (2003).

24. Fabrikant, G. et al. Computational model of membrane fission catalyzed by ESCRT-III. PLoS Comput Biol 5, e1000575 (2009).

25. Zhang, G. & Müller, M. Rupturing the hemi-fission intermediate in membrane fission under tension: Reaction coordinates, kinetic pathways, and free-energy barriers. J Chem Phys 147, 064906 (2017).

26. de Franceschi, N. et al. The ESCRT protein CHMP2B acts as a diffusion barrier on reconstituted membrane necks. J Cell Sci 132, (2018).

27. Bertin, A. et al. Human ESCRT-III polymers assemble on positively curved membranes and induce helical membrane tube formation. Nat Commun 11, 2663 (2020).

28. Pfitzner, A.-K. et al. An ESCRT-III Polymerization Sequence Drives Membrane Deformation and Fission. Cell 182, 1140–1155.e18 (2020).

29. Schöneberg, J. et al. ATP-dependent force generation and membrane scission by ESCRT-III and Vps4. Science 362, 1423–1428 (2018).

30. Remec Pavlin, M. & Hurley, J. H. The ESCRTs - converging on mechanism. J Cell Sci 133, (2020).

31. Johnson, D. S., Bleck, M. & Simon, S. M. Timing of ESCRT-III protein recruitment and membrane scission during HIV-1 assembly. Elife 7, (2018).

32. Abil, Z. & Danelon, C. Roadmap to Building a Cell: An Evolutionary Approach. Front Bioeng Biotechnol 8, 927 (2020).

33. van de Cauter, L. et al. Optimized cDICE for Efficient Reconstitution of Biological Systems in Giant Unilamellar Vesicles. ACS Synth Biol 10, 1690–1702 (2021).

34. Cada, A. K. et al. Friction-driven membrane scission by the human ESCRT-III proteins CHMP1B and IST1. Proc Natl Acad Sci U S A 119, e2204536119 (2022).

35. Roux, A., Uyhazi, K., Frost, A. & de Camilli, P. GTP-dependent twisting of dynamin implicates constriction and tension in membrane fission. Nature 441, 528–31 (2006).

36. Tucker, W. C., Weber, T. & Chapman, E. R. Reconstitution of Ca2+-regulated membrane fusion by synaptotagmin and SNAREs. Science 304, 435–8 (2004).

37. Ge, Y. et al. Two forms of Opa1 cooperate to complete fusion of the mitochondrial inner-membrane. Elife 9, (2020).

