## Supplementary figures and movie captions for "Dynamin A as a one-component division machinery for synthetic cells"

**Table of contents**

**Supplementary Figure 1:** Membrane topologies………………………………………………… 2
**Supplementary Note 1:** Explanation of quantification…………………………………………… 3
**Supplementary Figure 2:** Schematic depicting the expected values of $N_I$……………….....…… 5
**Supplementary Figure 3:** Median of recovery times in FRAP experiments…………………….. 6
**Supplementary Figure 4:** Open neck in the absence of DynA…………………………………... 7

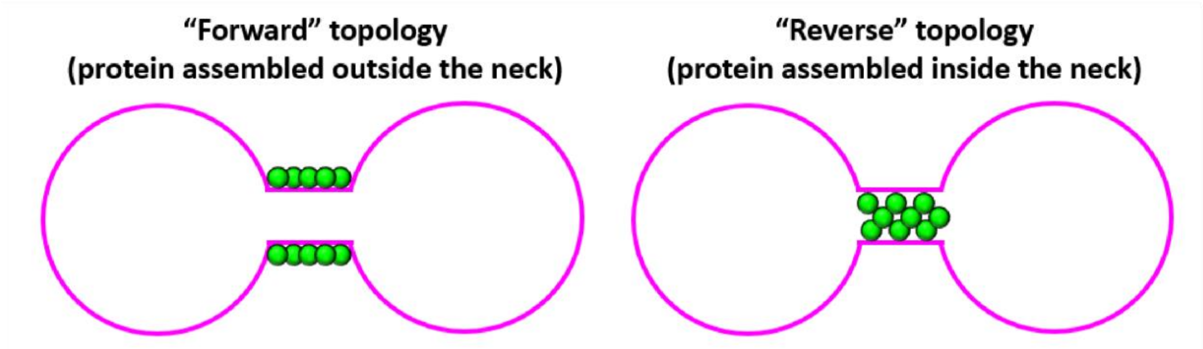

**Supplementary Figure 1: Membrane topologies**. Relative positions of membrane and protein machinery for the forward and reverse topology. Membrane is in magenta, protein in green.

### Supplementary Note 1: explanation of quantification

*Theoretical background*

In a typical FRAP experiment, one lobe of a multi-lobed chain of dumbbells is bleached and the recovery of fluorescent lipids across the neck is followed over time. The intensity of the bleached and control lobe are measured ($I_{FRAP}$ and $I_{CTRL}$, respectively), as well as the background intensity $I_{bckg}$. The background intensity is subtracted from the bleached and control lobe intensities. Furthermore, to account for residual fluorescence after bleaching, the intensity of the bleached lobe in the frame after the bleach, $I_{FRAP}(t = t_{FRAP})$ where $t_{FRAP}$ denotes the time point right after bleaching, is subtracted from both control and bleached lobe. Intensities corrected for background and residual fluorescence are denoted $\hat{I}$. The ratio between the intensities of the bleached and control lobe are computed before bleaching ($t \leq t_{FRAP} - \Delta t$) and used to normalize the recovery:

$$I_{norm}(t) = \frac{\hat{I}_{FRAP}(t)}{\hat{I}_{CTRL}(t)} / \frac{\langle \hat{I}_{FRAP}(t \leq t_{FRAP} - \Delta t) \rangle}{\langle \hat{I}_{CTRL}(t \leq t_{FRAP} - \Delta t) \rangle},$$

where $\langle \quad \rangle$ denotes a temporal average.

Only one control and one bleached lobes are quantified, irrespectively of the length and connectivity of the chain. The degree of recovery depends on the membrane connectivity at the neck: an open neck results in full recovery, a closed neck (full scission) results in no recovery, while hemi-scission leads to a partial recovery. Both full recovery and no recovery result in a well-defined value of $N_I$ (1 and 0, respectively, within the uncertainty due to the experimental error). In contrast, hemi-scission can result in values of $N_I$ that vary depending on the relative amount of membrane reservoir between the control and bleached lobes, which is the result of both size and number of lobes and their connectivity. This is explained in Supplementary Figure 2.

In the ideal case of an equal amount of membrane reservoir in both the control and bleached lobes, the value of $N_I$ is 0.33 (Figure S2a). If the membrane reservoir in the control lobe(s) exceeds the amount in the bleached lobe, $N_I$ increases up to a theoretical value of 0.5 for an infinite amount of membrane reservoir (Figure S2b). Conversely, if the amount of membrane reservoir is smaller in the control lobe(s), $N_I$ decreases, approaching a value of 0 for an infinitely large amount of membrane reservoir that is being bleached (Figure S2c). For this reason, the peak corresponding to hemi-scission in Figure 2b is relatively broad. It is not possible to normalize for the amount of membrane reservoir, because the connectivity between lobes cannot be established merely on the basis of imaging (i.e. the FRAP experiment itself is the only way to unambiguously determine the connectivity). Notably, there are also other experimental factors that can lead to broadening of the peaks, including lobes being slightly out of focus and residual fluorescence present in the bleached lobe after beaching.

*Explanation of automatic quantification*

Quantification of lobe intensities was performed in a semi-automated fashion. First, the centroid of the control and bleached lobes were roughly marked manually, after which two stages of automated refinement were implemented. First, edges in the fluorescence image were identified by a Canny edge detector (function *canny* from *scikit-image*) with $\sigma = 1$, followed by a Hough transform (function *hough_circle* from *scikit*-image) in order to identify circular objects, such as lobes. This results in a list of one to three candidate lobes, of which the one with its centre closest to the manually chosen lobe was selected as input to the second stage of refinement. The second stage of refinement consisted of a scan of lobe centers and lobe radii. Candidate radii were sampled uniformly between 5 and 15 pixels (corresponding to 1 to 3 μm) and candidate lobe centroids were sampled on a grid spanning 1/6 of the candidate radius found by the Hough transform around the user-marked centroid. This ensured sufficiently large sampling in both x-y location of the lobe centroid as well as its radius, yet restricted the search in order to avoid spurious results (such as the erroneous choice of a neighbouring lobe). For each combination of candidate centroid and radii, the fluorescence intensity was integrated by averaging over an annulus of 1.5 pixels width. The centroid and radius combination resulting in the maximum integrated intensity was chosen. For time trace quantification, the above scheme was applied to every frame of time lapse movies to a control lobe and the frapped lobe. The normalized FRAP trace was then computed the dividing the intensities of the FRAP-ped lobe by the intensities of the control lobe.

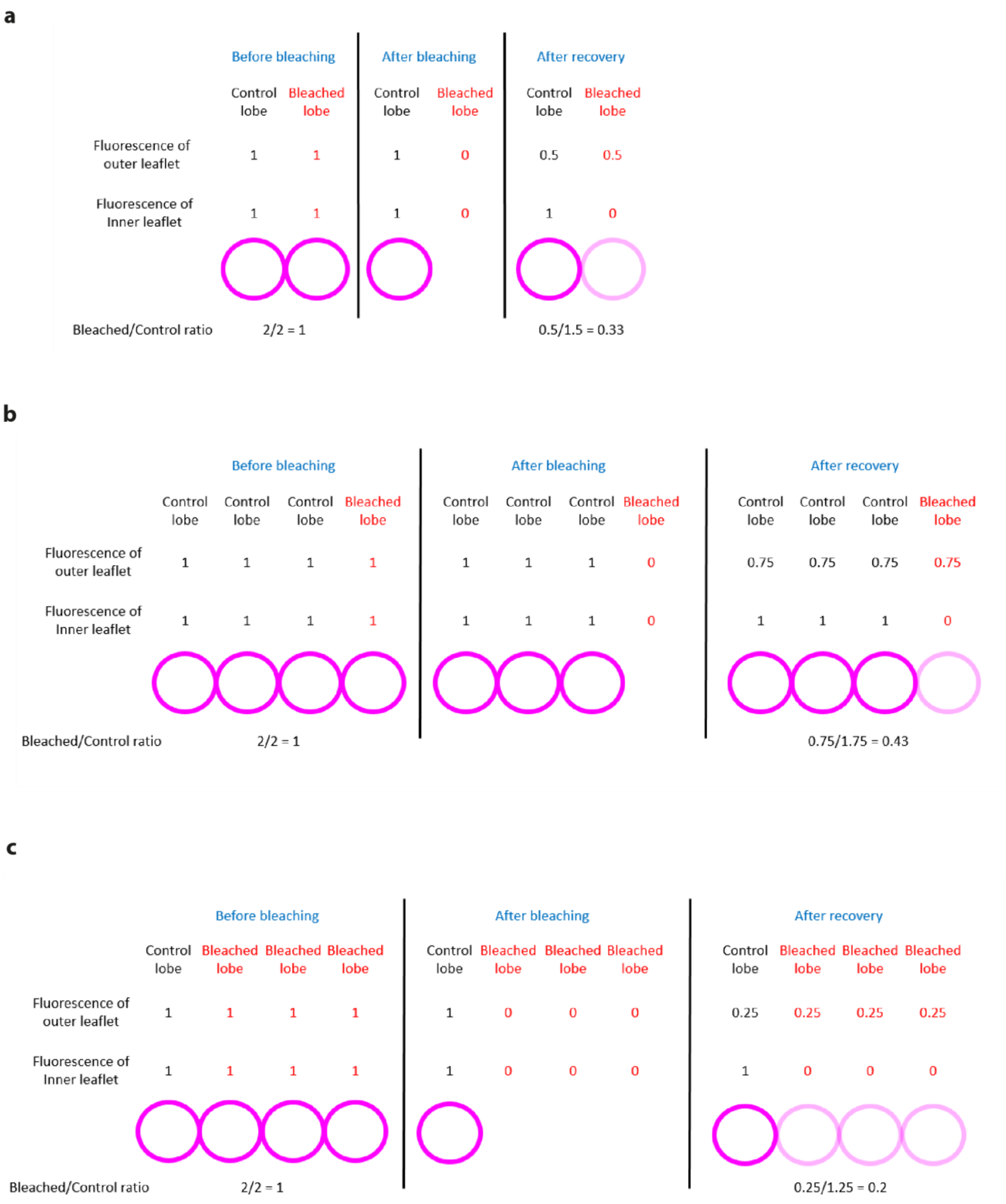

**Supplementary Figure 2:** Schematic depicting the expected values of $N_I$ for different amounts of membrane reservoir in control and bleached lobes**. a.** equal amount; **b.** larger reservoir in control lobe; **c.** larger reservoir in the bleached lobe.

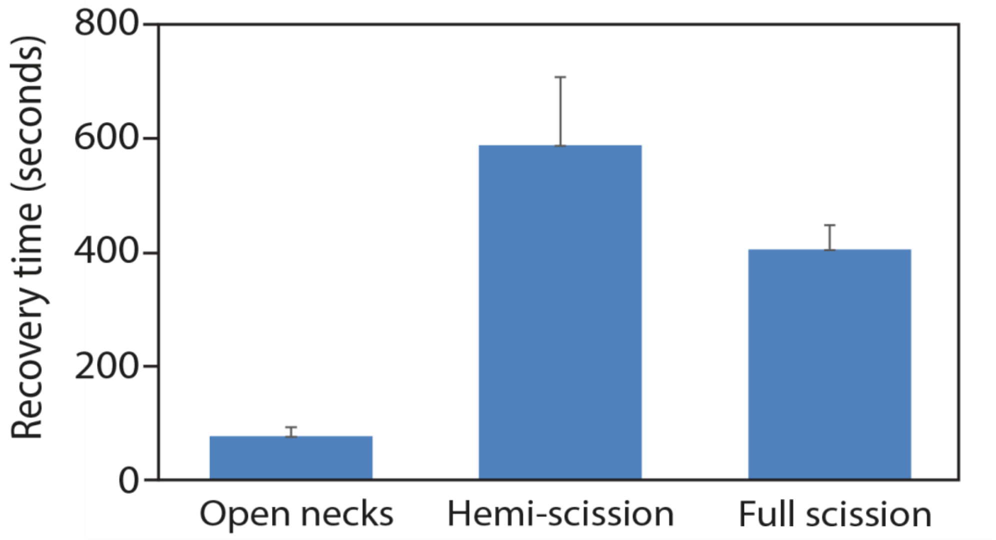

**Supplementary Figure 3:** Median of the duration of recovery times in FRAP experiments. Acquisition was stopped when no further lipid recovery was apparent. This quantitation is obtained from datasets such as shown in Figure 2a, 2b and S6b.

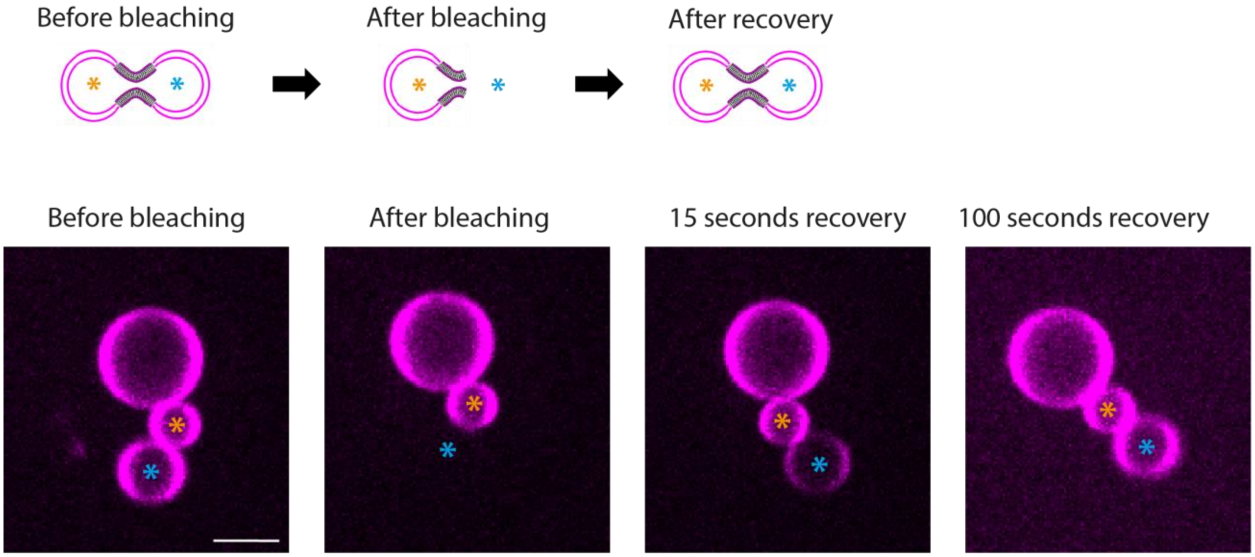

**Supplementary Figure 4: open neck in the absence of DynA.** Example of FRAP experiment showing full lipid recovery (open neck) on a chain of dumbbells without reconstitution of DynA. The orange asterisk indicates the control lobe, the blue asterisk indicates the bleached lobe. Scale bar: 5µm.

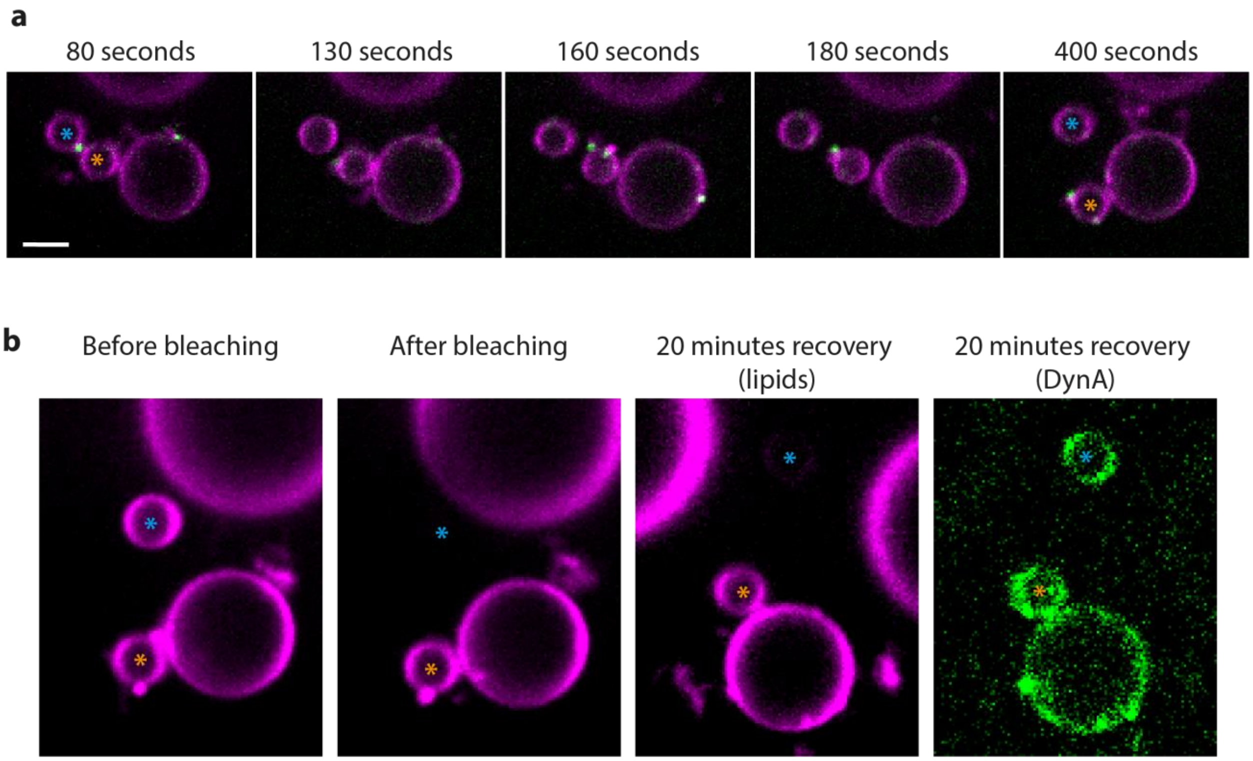

**Supplementary Figure 5: Lobe detachment upon scission.** (a) Confocal images of a dumbbell with a DynA cluster (in green) localized at neck. The dumbbell was imaged just as scission was occurring and the two lobes are drifting apart. (b) same dumbbell as in panel a. After lobes separation, FRAP analysis confirmed that the membrane connection between the two lobes had been lost and that no lipid recovery occurred. The bleached lobe is indicated by a blue asterisk, and its position is revealed by the low fluorescent signal of DynA binding to the membrane in the overexposed image.

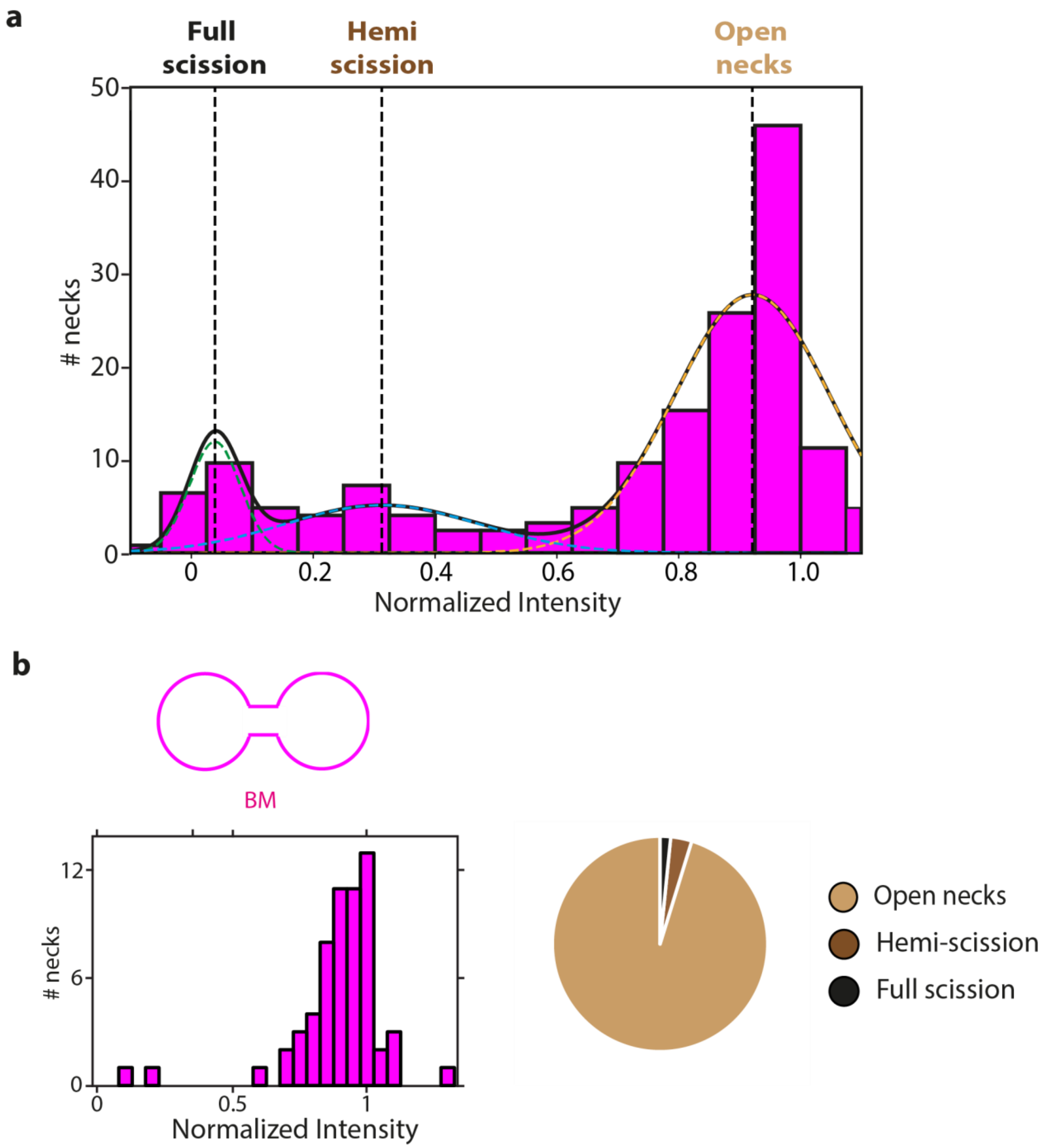

**Supplementary Figure 6: Quantification of the $N_I$ in different experimental conditions.**
**(a)** Quantification of the $N_I$ of a large dataset of necks from chains of dumbbells. This plot shows the data pooled from the three experimental conditions shown in Figure 2a, 2b and S6b (n=222 chains of dumbbells), and are decomposed into three peaks using a Gaussian Mixture Model. The three peaks correspond to open necks, hemi-scission and full scission. For each data point, the fluorescent lipids of one lobe were bleached and their recovery was followed over time until reaching the plateau. The final intensity was used for calculating and plotting the $N_I$. **(b)** Plot showing the $N_I$ of dumbbells having only bare membrane (left), and pie-chart (right) indicating the fraction of open necks, hemi-scission and full scission events (BN; 63 necks from 5 independent preparations).

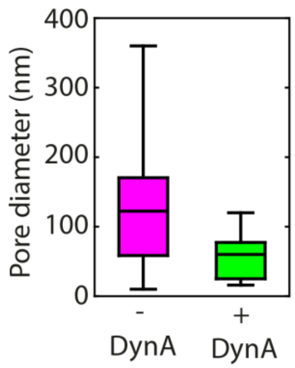

**Supplementary Figure 7: DynA assembly at the neck results in neck constriction.** Neck diameter in the presence or absence of DynA, estimated based on dye recovery upon photobleaching. In open necks and in the absence of DynA, the average inner diameter of necks was 134 ± 108 nm (mean ± SD). In the presence of DynA, the width of open necks appeared more constricted, with an average diameter of 57 ± 33 nm (mean ± SD). n=8 without DynA; n=9 with DynA. The estimation has been performed according to the procedure detailed in Ref. 14.

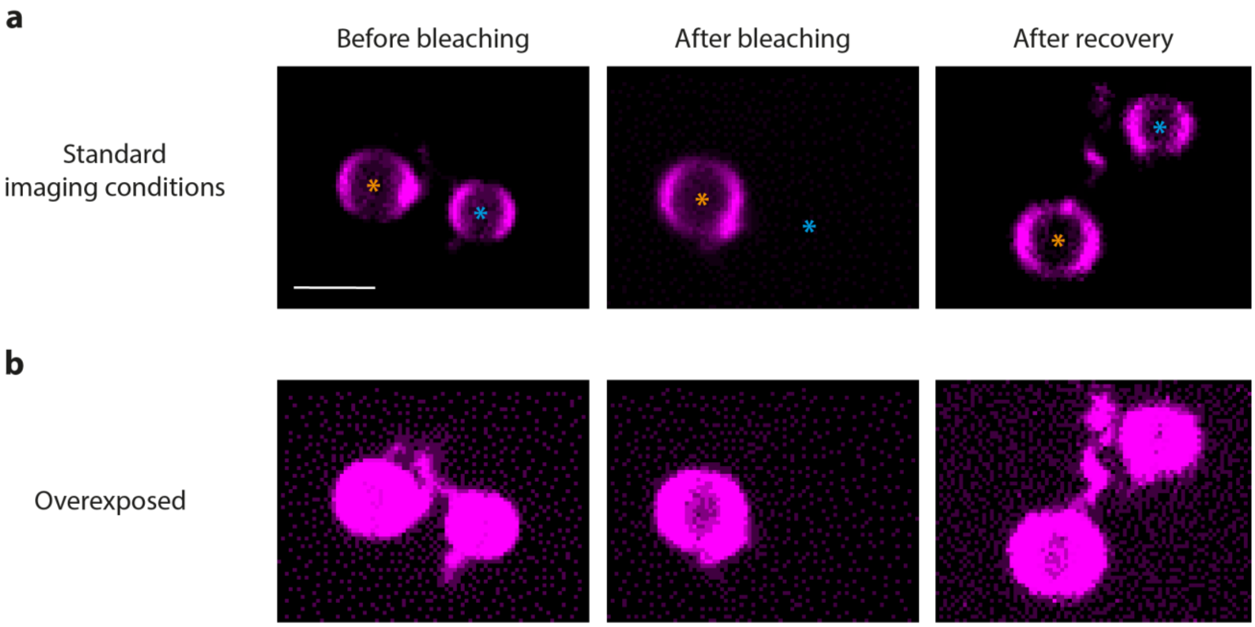

**Supplementary Figure 8: Liposomes connected by a nanotube.**

(a) Confocal image of two liposomes under standard imaging conditions which allow to clearly see the liposome membrane without overexposure. Upon bleaching of one liposome (marked with a blue asterisk) full lipid recovery could be observed. (b) Same images as in panel (a) with increased gain, causing overexposure of the liposome membrane. Such conditions reveal that a membrane nanotube is connecting the two liposomes and allows full lipid recovery.

**Movie captions**

**Movie 1: Stable localization of DynA at neck of a dumbbell liposome.** Movie of a confocal plane across a liposome. The movie shows a DynA cluster stably localized at the neck. Lipid fluorescence is shown in magenta, DynA in green. Frame rate was 1 image/sec.

**Movie 2: Liposome deformation into a chain of dumbbells.** Movie of a confocal plane across a liposome while it is being generated by the SMS approach. The movie captures the progressive deformation of the initially oblate liposomes into a chain of dumbbells. Liposome drifting while sinking is compensated by manually moving the stage. Lipid fluorescence is shown in magenta. Frame rate was 1 image/sec.

**Movie 3: Full recovery of fluorescent lipids in dumbbell liposome.** Movie of a confocal plane across a liposome during FRAP experiment. The movie shows lipid bleaching and progressive full recovery of fluorescent lipids flowing from the adjacent lobes through the neck. Lipid fluorescence is shown in magenta, DynA in green. Frame rate was 1 image/sec.

**Movie 4: Partial recovery of fluorescent lipids in dumbbell liposome.** Movie of a confocal plane across a liposome during FRAP experiment. The movie shows lipid bleaching and progressive partial recovery of fluorescent lipids flowing from the adjacent lobes through the neck. Lipid fluorescence is shown in magenta, DynA in green. Frame rate was 1 image/sec.

**Movie 5: Lack of recovery of fluorescent lipids in dumbbell liposome.** Movie of a confocal plane across a liposome during FRAP experiment. The movie shows lipid bleaching and the lack of recovery of fluorescent lipids. Lipid fluorescence is shown in magenta, DynA in green. Frame rate was 1 image/sec.
